# Bridging Worlds: Connecting Glycan Representations with Glycoinformatics via Universal Input and a Canonicalized Nomenclature

**DOI:** 10.1101/2025.05.30.657013

**Authors:** James Urban, Roman Joeres, Daniel Bojar

## Abstract

As the field of glycobiology has developed, so too have different glycan nomenclature systems, reflecting diverse cognitive and practical needs of different scientific uses. While each system serves specific purposes, this multiplicity creates challenges for usability, data integration, and knowledge sharing. Here, we present a practical framework for automated nomenclature conversion, taking any nomenclature as input, without having to declare the specific language, and using a canonicalized IUPAC-condensed format as a standardized output representation. Our implementation handles (i) all common nomenclatures, including common typos, (ii) complex cases including structural ambiguities, modifications, and uncertainty in linkage information, and (iii) different compositional representations. This Universal Input framework can translate more than 10 nomenclatures in less than 1 ms, tested on over 50,000 sequences with 95-100% coverage, enabling seamless integration of existing glycan databases and tools while maintaining the specific advantages of each representation system.

## Introduction

Glycans, complex carbohydrates attached to proteins and lipids, play crucial roles in numerous biological processes, including cell signaling, immune response, and disease progression (Varki 2017). As glycobiology has evolved over the past decades, so too have nomenclature systems used to represent these intricate molecules (Aoki-Kinoshita 2019). This diversity of representational systems reflects the varied needs of different scientific communities working with glycans, from carbohydrate chemists requiring precise atomic configurations, over glycoinformaticians working with unique and computationally tractable formats, to systems biologists analyzing glycome changes at scale.

While each nomenclature serves specific purposes, the resulting fragmentation creates significant challenges for data integration, knowledge sharing, and interdisciplinary collaboration. Experimentalists often favor nomenclatures optimized for human readability and manual annotation, such as the Oxford notation for *N*-glycans (Gornik et al. 2009) or IUPAC-condensed formats (McNaught 1996) that are frequently used (Jin et al. 2023). In contrast, computational glycobiologists often require machine-readable formats such as WURCS (Matsubara et al. 2017) or GlycoCT (Herget et al. 2008) that capture structural details with formal precision. Other nomenclatures, used in various niches of the glycosciences, include LinearCode^®^ (Banin et al. 2002), CSDB Linear (Toukach & Egorova 2020), or GlycoWorkbench (Ceroni et al. 2008). This divergence has created a gulf between manual experimental work and computational glycoinformatics that impedes the field’s advancement, both for information exchange and for accumulating large datasets required for AI models (Bojar & Lisacek 2022).

While individual converters exist (Klein & Zaia 2019; Tsuchiya et al. 2019), databases utilizing different nomenclature systems can remain siloed, software tools become nomenclature-specific, and researchers must master multiple representational languages to access the full breadth of glycobiology knowledge. This challenge is further compounded by inconsistent implementation of standards, typographical variations, and nomenclature dialects that have evolved within different research communities. This leads to a trend—unfortunately common in the glycosciences—in which each cluster of research groups independently, and repeatedly, develops a suite of methods that are hyper-optimized to their nomenclature preferences and data format peculiarities.

Previous efforts to address these challenges have typically proposed the adoption of a single standardized format, superior to previous formats. However, this approach overlooks the legitimate cognitive and practical advantages that different representations offer for specific use cases. IUPAC-condensed notation, for example, offers an intuitive balance between human readability, easy editability, and structural precision that makes it particularly valuable for manual annotation. Meanwhile, formats such as WURCS excel at representing structural ambiguities but present a steeper learning curve for human interpretation.

Rather than replace existing nomenclature systems, we propose to leverage their strengths through a Universal Input framework, capable of letting users work in any nomenclature they choose by automatically inferring any common nomenclature format, followed by its conversion into the widely used IUPAC-condensed format. Closely connected to this approach is the adoption of a canonicalized version of the IUPAC-condensed format, described here, as a standardized intermediate representation, exhibiting the best of both worlds. While standard IUPAC-condensed notation suffers from non-injectivity when mapping to other nomenclatures (i.e., several IUPAC-condensed strings can be mapped to the same WURCS glycan), and exhibits dialectical variations, these limitations can be easily and programmatically addressed through consistent canonicalization rules that we propose here, which take the burden of nomenclature conformity and validity from the user and place it on reactive and performant algorithms. We show further that this implementation is faster, more memory-efficient, and broader than current alternatives.

Our framework accommodates all common glycan nomenclatures, including typical typographical variations and errors, while handling complex cases such as structural ambiguities, modifications, and uncertainty in linkage information. Additionally, it processes compositional representations that specify monosaccharide content without fully defined structures, which presents a common output from mass spectrometry-based glycomics (Ruhaak et al. 2018).

By bridging the gap between manual experimental notation and computational glycoinformatics, this Universal Input framework enables researchers to work in their preferred nomenclature, while facilitating seamless data integration across the glycobiology ecosystem. Universal Input is implemented, and widely used, in the open-source Python library glycowork (Thomès et al. 2021), yet can be easily integrated into any Python-based application, such as our herein developed web application (https://canonicalize.streamlit.app/). The resulting interoperability gained from Universal Input promises to accelerate discovery by connecting previously isolated data repositories, enabling cross-platform tool development, and fostering interdisciplinary collaboration across the diverse communities studying glycan structure and function.

## Results

### Canonicalized IUPAC-condensed to balance computation and readability

While many glycan nomenclatures exist, we set out with the hypothesis that residue-focused nomenclatures offer the best balance between human-readability and computational processing (Fig. 1A). As outlined above, IUPAC-condensed is a human-readable notation that describes glycans while still being easily processable by computers (Fig. 1B), as we have shown repeatedly (Bennett et al. 2025; Bennett & Bojar 2024; Lundstrøm, Urban, & Bojar 2023; Thomès et al. 2023). The latter is also already amply demonstrated by the glycowork-internal parser (Thomès et al. 2021) and the grammar laying the foundation of the IUPAC-to-SMILES translator GlyLES (Joeres et al. 2023). Briefly, IUPAC-condensed uses common shorthand monosaccharide names, where only the uncommon form is further specified (e.g., Glc instead of D-Glcp), which are connected via parenthesis-enclosed linkages containing (i) the anomeric configuration, (ii) the anomeric carbon, and (iii) the recipient hydroxyl group on the subsequent monosaccharide, such as in Gal(b1-4)Glc. Branches are enclosed in square brackets, which can be nested, while residues with uncertain connection to the remaining glycan are enclosed in curly brackets. Modifications are typically listed as a number (indicating the carbon at which this modification is found), followed by a shorthand notation of the modification (e.g., Gal6S). Ambiguities can be further expressed on the monosaccharide level (e.g., Hex instead of Gal) and on the linkage level (e.g., a1-?, ?1-4, or the more granular a1-3/4).

**Figure 1.**
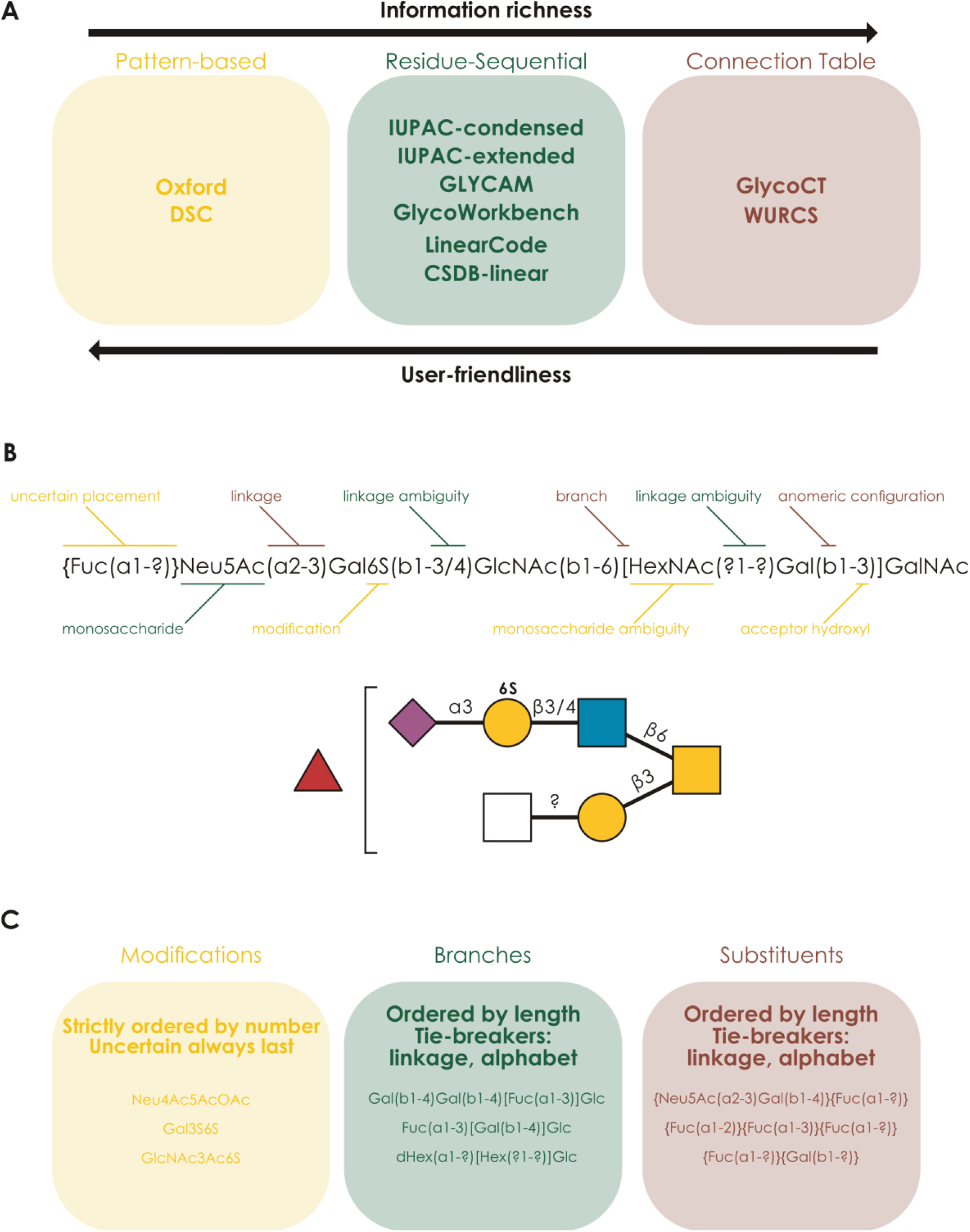
IUPAC-condensed as a balanced nomenclature for glycobiology. **A)** An overview of common glycan nomenclatures and their properties. **B)** Anatomy of the IUPAC-condensed nomenclature, with the example of an *O*-glycan, visualized in the Symbol Nomenclature For Glycans (SNFG) via GlycoDraw (Lundstrøm, Urban, Thomès, et al. 2023). **C)** Ordering guidelines for converting a regular IUPAC-condensed glycan into canonicalized IUPAC-condensed, as defined here. We emphasize that we do not intend this as a user manual but rather a transparent explanation of the ordering guidelines employed by our canonicalization algorithms.

One key property of the classic IUPAC-condensed notation is that multiple semantically valid notations exist for a single glycan, e.g., Gal(b1-3)GalNAc(b1-4)[Neu5Ac(a2-3)]Gal(b1-4)Glc and Neu5Ac(a2-3)[Gal(b1-3)GalNAc(b1-4)]Gal(b1-4)Glc. There are also countless dialects and personal notational preferences (the forgivingness for such undoubtedly contributing to the nomenclature’s popularity among experimentalists) that need to be considered. To simplify working with IUPAC-condensed computationally, we thus set out to canonicalize its notation, defining one unique description per glycan. We emphasize here that these canonicalizations are not meant as a prescriptive burden to users of IUPAC-condensed. Rather, they are intended as algorithmically enforceable routines to surjectively convert any arbitrary IUPAC-condensed starting point into a unique, canonicalized string (that is still human-readable and editable). The canonicalization of IUPAC-condensed consists of two main parts: (i) enforcing “standard” IUPAC-condensed (i.e., remedying dialects and typos) and (ii) ordering modifications, floating substituents, and branches in a deterministic manner (Fig. 1C).

While Gal(b1-4)Glc, Galb1-4Glc, and Galb4Glc all describe lactose, only the first one would be considered standard IUPAC-condensed. Within glycowork, and in the *canonicalize_iupac* function described and discussed below, we employ extensive regular expression operations to guarantee the proper namespace of monosaccharides, modifications, linkages, as well as many other variations. While we discuss other nomenclatures below, this already allowed us to homogenize the great diversity within the realm of IUPAC-condensed, converting them into standard IUPAC-condensed nomenclature.

Next to following these conventional IUPAC-condensed guidelines (McNaught 1996), our canonicalization algorithms (see Methods) also follow strict ordering guidelines that circumvent the inherent ambiguity of the nomenclature. We here emphasize that the particular choice of ordering is almost irrelevant, and the main benefit derives from having guidelines that can be (algorithmically) enforced from any starting point. With the example of branch ordering (Fig. 1C), we employ a recursive, graph-based, approach that iteratively designates the longest (i.e., most monosaccharides) branch as the main chain, with several tiebreakers (e.g., linkage at the branchpoint) that ensure a unique ordering, which, on average, minimizes nested branches.

For the sake of consistency and the ease of writing deterministic/efficient code, we elect to omit the reducing end ‘dangling’ linkage (e.g., ‘GlcNAc(b1-’) from canonicalized IUPAC-condensed. Yet we argue that this information is almost entirely implicit in any case, since (i) protein-linked glycans are invariant in this aspect (i.e., *N*-glycans always have beta-linked GlcNAc at their reducing end), (ii) repeating units contain this information explicitly due to our convention of writing them as wrap-arounds, and (iii) free glycans (e.g., milk glycans) cycle between alpha and beta in solution.

We note that the convenience of having a guaranteeable and regular nomenclature format has allowed us to implement the extremely efficient glycan processing operations within glycowork, such as crafting a glycan-specific regular expression system (Bennett & Bojar 2024), and we are convinced this set-up will spur more advances in the glycosciences.

### The Universal Input system infers and auto-converts all common nomenclatures

The main issue with proposing new nomenclature variants inevitably is user conformity (and, hence, validity of the canonicalized nomenclature). We propose to solve this issue here with our main advance of this work, the Universal Input system (Fig. 2), in which any deviation from this canonicalized nomenclature (e.g., different nomenclatures, nomenclature dialects, arbitrary branch ordering, typos) is automatically fixed, such that users do not need to comply with (or even be aware of) syntactic rules, as they can be algorithmically enforced from any common starting point, without appreciable computational overhead as we show below.

**Figure 2.**
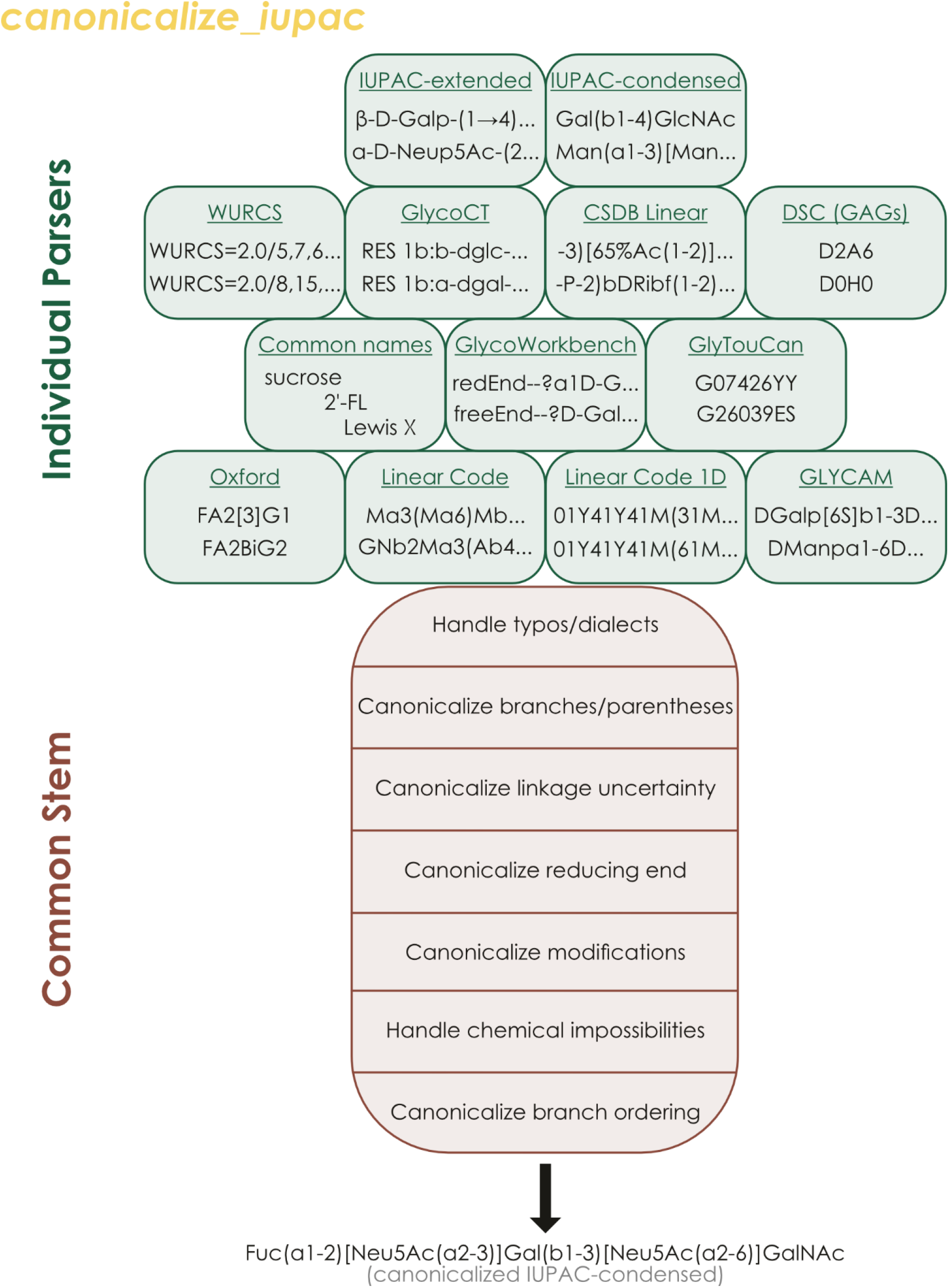
Universal Input is a potent platform to enable users to work in any common glycan nomenclature. Glycan sequences used as input for *glycowork.motif.processing.canonicalize_iupac* first are searched for unique hooks indicating a certain nomenclature, after which they are dispatched to the dedicated parser for that nomenclature, followed by all sequences passing through the common stem, in which dialectical variation in syntax, monosaccharide name space, modification handling, and branch ordering are sanitized and canonicalized to arrive at a unique canonicalized IUPAC-condensed string per molecule.

In general, such a system requires two operations: (i) a robust means to detect a deviation and (ii) an algorithmic solution to remedy said deviation. We distinguish here between different nomenclatures and notational variance of IUPAC-derived and similar nomenclatures. For dedicated nomenclatures (Fig. 2), we identified unique string- or regular expression-patterns (‘hooks’; see Supplementary Table 1) that allowed us to call a specific parser for that nomenclature, which we newly implemented for this work. Generally, the consolidation of ‘post-processing’ operations into a common stem has allowed us to keep parsers relatively lightweight and fast to construct, as their task was merely to convert sequences into anything resembling IUPAC nomenclature, without strict requirements on particular notation, making this system easily extendable to new nomenclatures.

We note that, tested on >150,000 sequences (Supplementary Table 2; see also Fig. 3 below), our hooks for nomenclature detection are empirically robust to typically used sequences and all common nomenclatures. The common stem, after the parsers, then mainly used carefully crafted regular expressions for many common deviations (e.g., in monosaccharide namespace, linkage notation, modification usage, etc.), collected and refined over the past three years. Lastly, branching order was algorithmically canonicalized, based on the rules mentioned above, resulting in a unique string per molecule that is (i) human-readable, (ii) easily editable, and (iii) amenable to be analyzed by performant analyses such as presented within glycowork. This entire workflow is consolidated in the single *canonicalize_iupac* function within glycowork (as well as in easily exportable decorators, as mentioned below). Even more importantly, this function (and its connected decorator) is widely used in glycowork itself, making the package compatible with all common nomenclatures, and, e.g., allows users to draw SNFG-depictions of glycans by calling the GlycoDraw function on WURCS sequences. We emphasize that *canonicalize_iupac* (and thus Universal Input) does not require any other information than a string-based glycan sequence, as everything else will be automatically inferred, making it very user-friendly and robust.

**Figure 3.**
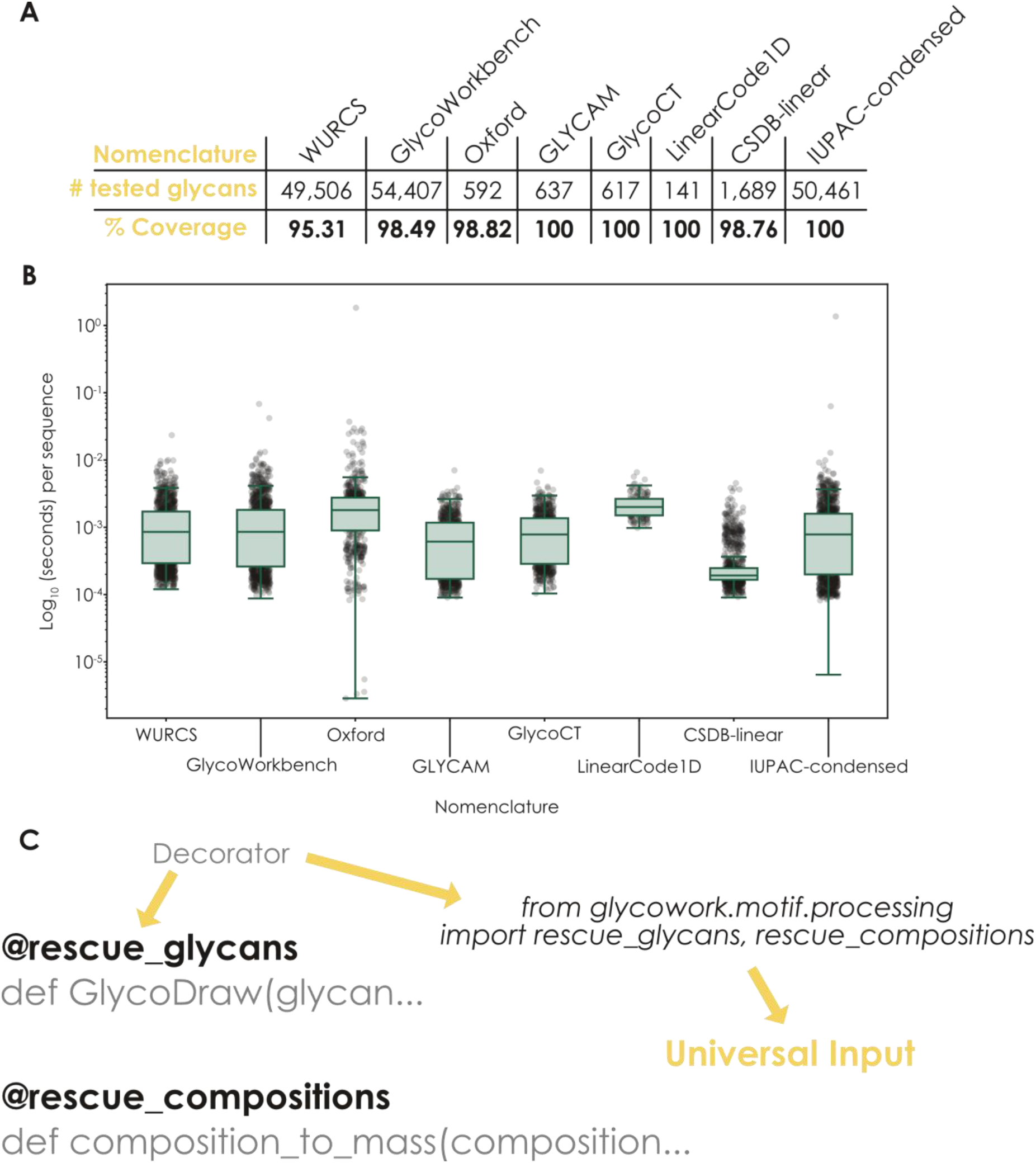
Universal Input is robust, performant, and readily usable anywhere in Python. **A)** Coverage of common nomenclatures by Universal Input. For the glycan nomenclatures of WURCS, GlycoWorkbench, Oxford, GLYCAM, GlycoCT, LinearCode, CSDB-linear, and IUPAC-condensed, we have assembled a set of representative sequences (Supplementary Table 2) and tested them with the *canonicalize_iupac* function, shown here as the success rate of this function. **B)** Speed test of Universal Input. For the sequences mentioned in (A), we then also timed the execution speed of the *canonicalize_iupac* function per nomenclature, visualized as box plots (the line being the median values, with box edges indicating quartiles, and whiskers indicating the remaining data distribution up to the 95% confidence interval), overlaid with the results of up to 1,000 randomly chosen sequences as a swarm plot. We report average runtimes of 1 ms per glycan, when run on a default Google Colaboratory instance (Intel^®^ Xeon^®^ CPU @ 2.20GHz). **C)** Enabling Universal Input in any Python code. Using the Universal Input decorators for sequences (*rescue_glycans*) and compositions (*rescue_compositions*), functions in Python can be equipped with Universal Input capabilities with a single line of code, imported from glycowork.

Further, while the line between sequences and compositions can be blurry and *canonicalize_iupac* supports some structured compositions such as “(Hex)3 (HexNAc)1 (NeuAc)1 + (Man)3(GlcNAc)2”, we also developed the *canonicalize_composition* function to support the great diversity of composition formats used in the academic literature (e.g., H5N4F1A2 or Hex5HexNAc4Fuc1Neu5Ac2; Supplementary Fig. 1), which we also use within glycowork to support mass calculations and other operations with any common composition nomenclature.

We next set out to establish the coverage and robustness of our Universal Input system by testing the *canonicalize_iupac* function on up to 50,000 sequences per nomenclature (Supplementary Table 2), resulting in coverage values ranging from 95-100% (Fig. 3A). Since the Universal Input functions often run in the background for operations within glycowork, we needed to ensure that they still stayed as performant as possible for a seamless user experience. We thus benchmarked the speed of *canonicalize_iupac* on the sequences used for our coverage test above and, despite numerous regular expression operations and other procedures, can report an average processing time of less than 1 ms per glycan on regular consumer hardware (Fig. 3B), regardless of nomenclature. We also compared *canonicalize_iupac* with the format conversion within glypy (Klein & Zaia 2019) on WURCS, GlycoCT, and LinearCode^®^ (since other nomenclatures were not supported on glypy) and report that, on average, Universal Input was 1.5-2x faster and used 40-60% less memory than glypy (Supplementary Fig. 2A). Further, for usage in Python our Universal Input platform was faster than converting sequences via the GlycanFormatConverter API (Supplementary Fig. 2B), and offered more features than both alternatives (Supplementary Fig. 2C).

Lastly, while we both offer Universal Input to users and elevate glycowork’s functionality with it, we reasoned that it might be modular enough to be integrated into any Python package related to glycobiology. For this, we offer two options: (i) importing the actual *canonicalize_iupac* / *canonicalize_composition* functions, for granular functional control over where exactly canonicalization should occur, or (ii) the new *rescue_glycans* / *rescue_compositions* decorators. These decorators can be prepended before every function that uses glycan strings (or lists of strings) as input and then automatically equip that function with Universal Input capabilities (Fig. 3C). We have tested this system recently in our CandyCrunch (Urban et al. 2024) and GlyContact (https://github.com/lthomes/glycontact) packages and achieved Universal Input of each package with just a few lines of code, with full control over which functions should and should not benefit from this system. We thus conclude this section by arguing that Universal Input makes the glycowork package an even more attractive starting point for glycoinformatics, both in itself, as well as in building capacity for constructing downstream Python applications.

### Making Universal Input accessible yields new opportunities for glycobiology

We note that, since *canonicalize_iupac* operates on a per-glycan level, there is also no need for users to exclusively use one nomenclature. Datasets with any mixture of nomenclatures can be readily used for any workflow within glycowork (Supplementary Fig. 3). We can envision at least two use-cases for which this is useful: (i) any comparative analysis endeavor, up to a full meta-analysis, will have gathered data in all kinds of nomenclatures, which is easily handled with this functionality, and (ii) datasets containing one or a few glycans with not (yet) supported sequence features, such as unusual modifications, can still largely be analyzed with most of the functionality within glycowork.

We realized that this platform of not only converting nomenclatures but also canonicalizing conventional nomenclatures, as well as cleaning them from typos and other errors, could be generally useful for the broader audience of non-Python glycobiologists. Therefore, we have exposed the *canonicalize_iupac* functionality via our no-code graphical user interface (GUI) for glycowork that is freely available for Windows and Mac via https://github.com/BojarLab/glycowork/releases (Fig. 4A).

**Figure 4.**
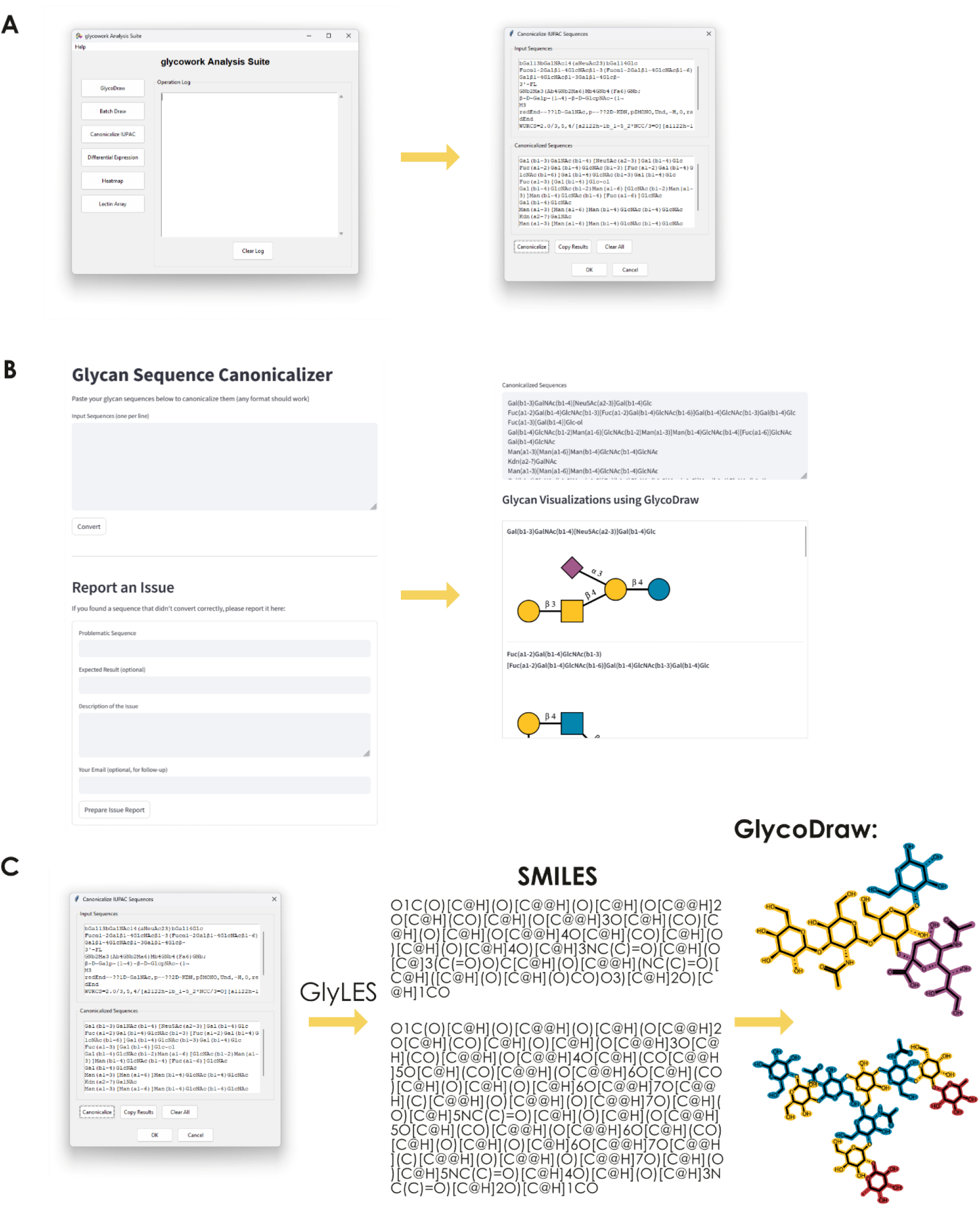
Universal Input is easily accessible and unlocks access to chemoinformatics. **A)** Using Universal Input via the glycoworkGUI (v1.6.1). The graphical user interface of glycowork exposes the *canonicalize_iupac* function as “Canonicalize IUPAC”, which allows users to input a number of sequences in any nomenclature format and copy the output into anywhere they choose. **B)** Using Universal Input via our web application. On https://canonicalize.streamlit.app, users can input any number of glycan sequences in any format, convert them into canonicalized IUPAC-condensed with a single click, and copy them or view them via GlycoDraw-drawn SNFG depictions in a scroll box. **C)** Using Universal Input to convert any glycan nomenclature into SMILES strings. Any canonicalized IUPAC-condensed sequence obtained from our Universal Input platform can be converted into SMILES strings using the *glycowork.motif.processing.IUPAC_to_SMILES* function (which relies on the GlyLES converter we previously developed). This then interfaces with existing chemoinformatic functionality, such as drawing glycans on an atomic level using GlycoDraw within glycowork.

To expand the reach of our Universal Input platform even further, we then also developed a web application using Streamlit that allowed users to rapidly copy/paste their to-be-converted sequences in plain text (https://canonicalize.streamlit.app/; Fig. 4B). For maximum utility, this web-app also used our GlycoDraw (Lundstrøm, Urban, Thomès, et al. 2023) platform to visualize converted sequences in a SNFG format, allowing users to quickly assess whether the conversion worked as expected (or export SNFG drawings), even if they are unfamiliar with the canonicalized IUPAC-condensed nomenclature. We also added an error report form to this application, facilitating interaction with the community and further improvement of the Universal Input platform.

Lastly, we wanted to stress the lingua franca aspects of our canonicalized IUPAC-condensed approach by showcasing the connection to state-of-the-art glycoinformatics capabilities this facilitates. One example of this can be shown via the integration with the GlyLES platform (Joeres et al. 2023), which converts IUPAC-condensed glycans into SMILES strings. These well-established chemical descriptions of molecules are not only opening doors to the entirety of chemoinformatics (RDKit etc.) but also directly enable mass calculations and other essential operations to characterize glycans. *De facto*, our Universal Input platform makes every glycan nomenclature easily convertible into SMILES strings (without the need for dedicated GlycoCT-to-SMILES parsers etc.), by first converting it into canonicalized IUPAC-condensed as a ‘pivot language’ that is then compatible with approaches such as GlyLES (Fig. 4C). To demonstrate this, we have used this to calculate and compare the normalized topological polar surface area (TPSA) for glycans in three databases (glycowork, GlyCosmos, GlycoWorkbench), written in three different nomenclatures (IUPAC-condensed, WURCS, GlycoWorkbench), demonstrating the potential of this approach.

## Discussion

The Universal Input framework presented here addresses several critical challenges in glycoinformatics while offering distinct advantages over existing approaches. One yet unmentioned significant benefit is the offline functionality of our framework, enabling nomenclature conversion without requiring internet connectivity or external service dependencies. This feature ensures accessibility in diverse research environments and delivers superior performance, with conversion speeds orders of magnitude faster than web-based alternatives (Supplementary Fig. 2B).

Our approach also lays the groundwork for improved biocuration practices in glycobiology (Martinez et al. 2024). By providing a standardized method for nomenclature conversion, this framework enables more consistent annotation of glycan structures across databases and literature. Researchers can leverage the canonicalized IUPAC-condensed format as a reference representation while maintaining compatibility with their preferred nomenclature systems. Automated data mining approaches (Kahsay et al. 2025) could use Universal Input to accommodate common variations in glycan notations during data extraction. This standardization facilitates developing quality control mechanisms for detecting and correcting inconsistencies in glycan structure databases. Overall, we emphasize that our main goal is not to derive a theoretically perfect nomenclature, but to craft a system that is maximally useful in real-world practice, by taking the burden of nomenclature validity from the user.

The modular architecture of our framework (rudimentary parsers + a common stem) represents another key advantage. Adding support for a new nomenclature system requires only the development of a rudimentary parser that converts the format to anything remotely close to IUPAC-condensed notation—often only a few lines of code—along with a “unique hook” detection mechanism that identifies the nomenclature from input strings. If no sufficiently performant hook can be identified, this is usually an indicator of strong IUPAC-condensed resemblance (e.g., CSDB-Linear) and can be handled by the common stem. This extensibility ensures that the framework can evolve alongside the field, accommodating both established and emerging nomenclature systems. Our modular design also allows for targeted improvements to specific parsers without disrupting the overall conversion pipeline, facilitating continuous refinement made possible by our open-source software.

We argue that the canonicalized IUPAC-condensed format serves as an effective lingua franca for glycan representation, balancing human readability with computational precision. While the original IUPAC-condensed notation suffers from ambiguities and dialectical variations, our canonicalization rules establish a consistent interpretation that preserves intuitive structure, while enabling reliable computational processing. This standardized representation provides a foundation for powerful back-end applications, such as the glycan graphs used in glycowork (Thomès et al. 2021).

Despite these advances, several challenges remain. The complex and often inconsistent nature of glycan nomenclature systems means that edge cases will inevitably arise that require refinement of our conversion algorithms, which is why the open-source nature of our code is so crucial. The open-source model enables specialists across the entire field to contribute improvements, extending the coverage and accuracy of this framework. Community-driven development also helps distribute the maintenance burden, increasing the likelihood that Universal Input will remain viable as glycoinformatics continues to evolve.

Additionally, while our framework handles the syntactic aspects of nomenclature conversion effectively, addressing the semantic layer (i.e., ensuring that converted structures accurately represent the intended molecular reality) requires both active curation and ongoing validation against experimental data. Future work should focus on expanding our coverage of rare monosaccharides and unusual modifications in various nomenclatures while developing more sophisticated validation mechanisms to assess conversion accuracy or capture unphysiological structures.

## Methods

### Universal Input

Within the Universal Input system, a combination of dedicated parsers and a common stem comprise the *canonicalize_iupac* function. We decompose this into several blocks, further described below: (i) nomenclature detection via hooks, (ii) individual parsers into basic IUPAC-condensed, (iii) a common stem to streamline IUPAC-condensed variations and general clean-up, and (iv) a branch canonicalization algorithm to yield canonicalized IUPAC-condensed sequences. The Universal Input system, as described here, is available in glycowork (v1.6.1+), the glycoworkGUI (v1.6.1+), and a dedicated web interface (https://canonicalize.streamlit.app/). If used via Python, Python >= 3.9 is required.

### Nomenclature detection

First, common names (e.g., “LacNAc”) are tried to be accessed in a glycowork-stored dictionary with a runtime complexity of O(1), for which we ignore capitalization and spaces. Next, the presence of specific hooks in the glycan sequence (see Supplementary Table 1) will trigger the *linearcode_to_iupac, linearcode1d_to_iupac, iupac_extended_to_condensed, glycoct_to_iupac, wurcs_to_iupac, glycam_to_iupac, GAG_disaccharide_to_iupac, glytoucan_to_glycan, nglycan_stub_to_iupac*, or *oxford_to_iupac* parser in *glycowork.motif.processing*. The usage of yet-unsupported nomenclatures, such as SMILES, is detected via the *check_nomenclature* function and will lead to an appropriate error.

### Individual parsers

All parsers heavily use regular expressions and other code operations to convert the original nomenclature into a flat string resembling IUPAC-condensed nomenclature. WURCS, GlycoCT, and, to some extent, Oxford are first parsed into dictionaries of connections and building blocks, which are then used to construct the final sequence. WURCS parsing in particular relies on a mapping of valid tokens, stored within glycowork. We stress that the individual parsers are not meant to be used in isolation because, by design, they do not result in clean and final IUPAC-condensed nomenclature. For this, we use the common stem of *canonicalize_iupac*, described below, to perform any shared operations (such as canonicalizing the usage of parentheses and brackets etc.).

### Cleaning IUPAC-condensed sequences in a common stem

In general, the common stem of *canonicalize_iupac* uses extensive regular expressions to capture, homogenize, and standardize nomenclature syntax. As a first step, common token variants are detected and replaced (e.g., NeuAc → Neu5Ac or β → b). Other operations here include sanitizing variations in the placement of anomeric and enantiomeric indicators, in the usage of dashes or not in linkages, in the convention of explicitly declaring the starting carbon or not, and many others. Another common theme here is the standardization of linkage ambiguity (e.g., occasionally noted as: Mana-, Man-, Mana1-, Man[, Man-[, ManMan, and many other variations). Next, we standardize the usage of parentheses vs brackets. If parentheses are used for branches, this is changed to the conventional style of parenthesis = linkage, bracket = branch. Similarly, we standardize the usage of glycan modifications, where we encountered (and fixed) variants such as: [4S]Gal, Gal?S, [S]Gal, SGal, S-Gal, and many more. Next, variants of denoting uncertain substituents (i.e., “we know there is a sialic acid in this glycan but not where”) are standardized to the format of curly bracket = floating substituent. Thereafter, the common stem automatically checks for chemical impossibilities (e.g., linkages where no acceptor hydroxyl group is available or multiple linkages ending in the same acceptor hydroxyl group) and sets the corresponding linkages to ?-containing wildcards that resolve the impossibility. We note that this common stem also converts CSDB-Linear into IUPAC-condensed, both because we did not find a unique hook for a CSDB-specific parser but also because many of the operations are shared between the two nomenclatures.

### Canonicalization of branch ordering

If the used glycan sequence at the end of the common stem contained branches, we used a branch canonicalization algorithm that resulted in a unique IUPAC-condensed sequence per glycan. This is done by converting the glycan into a NetworkX-based directed graph (via *glycowork.motif.graph.glycan_to_nxGraph*), reordering branches on the graph level, and then translating that graph back into a canonicalized IUPAC-condensed sequence via the *glycowork.motif.graph.graph_to_string* function (using the *canonicalize_glycan_graph* function). For the actual branch canonicalization, we use a recursive depth-first traversal approach. First, it calculates values using post-order traversal (where children are processed before parents), and then builds the canonical graph using pre-order traversal (where parents are processed before children). The sorting of branches happens at each node during this depth-first process, prioritizing the longest paths (i.e., the branch with the longest chain of monosaccharides, recursively). When two branches have equal path lengths, tie-breaking is performed first by the linkage number of the branch parent (lower numbers are preferred; e.g., Man(a1-3) branch before Man(a1-6) branch, in *N*-glycans), with integer linkages prioritized over wildcards. If linkage numbers are also equal, branches are further distinguished alphabetically by comparing the minimum leaf labels (or proceeding down the branch until an alphabetic difference is detected), ensuring a completely deterministic ordering.

### Collecting glycan sequences for benchmarks

For WURCS sequences, we gathered all sequences labeled “Linkage-defined saccharide” that were available on GlyCosmos (Yamada et al. 2020) (April 2025). For GlycoWorkbench, we gathered all available sequences from https://gitlab.com/glycoinfo/glycoworkbench. For Oxford, we gathered representative examples from the following GlycoPOST IDs (Watanabe et al. 2021): GPST000308, GPST000318, GPST000421, GPST000473, GPST000494. For GLYCAM and GlycoCT, we gathered all sequences that were available on GlycoShape (Ives et al. 2024) (May 2025). For LinearCode1D, we gathered sequences from various GlycoPOST IDs. For CSDB-Linear, we were supplied with a list of representative sequences from Dr. Tom Stanton (https://github.com/BojarLab/glycowork/issues/76). For IUPAC-condensed, we gathered all sequences in the *df_glycan* dataset in glycowork (v1.6.1) (Thomès et al. 2021).

## Code availability

The Universal Input framework is available, actively used by, and accessible from glycowork (https://github.com/BojarLab/glycowork), v1.6.1+.

## Acknowledgment

This work was funded by a Branco Weiss Fellowship – Society in Science awarded to D.B., by the Knut and Alice Wallenberg Foundation, and the University of Gothenburg, Sweden. The authors would like to thank Kiyoko Aoki-Kinoshita, Frédérique Lisacek, Thomas Lütteke, Sriram Neelamegham, and Issaku Yamada for fruitful discussions about the IUPAC-condensed notation.

## Contributions

D.B. conceptualization; D.B., J.U., R.J. formal analysis; D.B. resources; D.B., J.U., R.J. data curation; D.B., J.U., R.J. writing–original draft; D.B., J.U., R.J. writing–review & editing; D.B., J.U. visualization; D.B. supervision; D.B. funding acquisition; D.B., J.U., R.J. methodology; D.B., J.U., R.J. validation.

