## Supplemental Figures for "Bridging Worlds: Connecting Glycan Representations with Glycoinformatics via Universal Input and a Canonicalized Nomenclature"

### Supplementary Figures

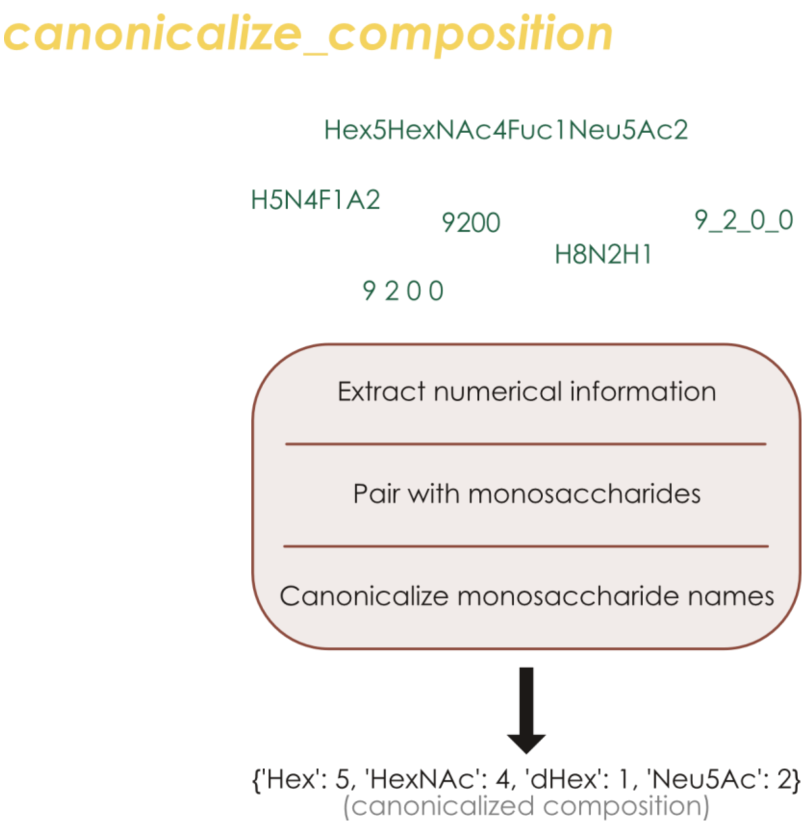


**Supplementary Figure 1. Universal Input supports various composition notations.** Examples of supported composition formats are shown, together with the processing steps performed in *glycowork.motif.processing.canonicalize_composition*, leading to a standardized dictionary of type monosaccharide : quantity. We note that the monosaccharide namespace of the output is controlled and dictionaries are insertion-ordered to always result in the exact same dictionary for two glycans with identical compositions, allowing us to hash it for fast comparisons.

**A**

| Nomenclature | WURCS | GlycoCT | LinearCode | |
|---|---|---|---|---|
| Speed improvement | +109% | +157% | +205% | higher is better |
| Memory improvement | -44% | -39% | -62% | lower is better |

**B**

| Universal Input | Glycan Format Converter |
|---|---|
| 1k WURCS Timing | |
| 0.4s | 910s |
| 1k WURCS Accuracy | |
| 99.2% | 100% |
| 100 GlycoCT Timing | |
| 0.012s | 244s |
| 100 GlycoCT Accuracy | |
| 95% | 91% |

**C**

| | Glycan Format Converter | Universal Input | glypy |
|---|---|---|---|
| Input languages | 7 | 12 | 5 |
| Handles structural ambiguity | Yes | Yes | Yes |
| Automatic language detection | No | Yes | No |
| Accessibility | Web interface, API, Java code | Web interface, GUI, Python code | Python code |

**Supplementary Figure 2. Universal Input is faster and more efficient than alternatives. A)** Comparing our benchmark sequences for WURCS, GlycoCT, and LinearCode® with the *glypy.io* module (v1.0.17) and the *canonicalize_iupac* from glycowork (v1.6.1) revealed substantial improvements in speed and memory usage, when using Universal Input. **B)** Comparing 1,000 WURCS sequences and 100 GlycoCT sequences with *canonicalize_iupac* from glycowork (v1.6.1) and the GlycanFormatConverter API demonstrated orders-of-magnitude speed improvements via the Universal Input platform within Python, while maintaining acceptable accuracy. **C)** Feature comparison of the nomenclature conversion systems relevant for usage in Python.

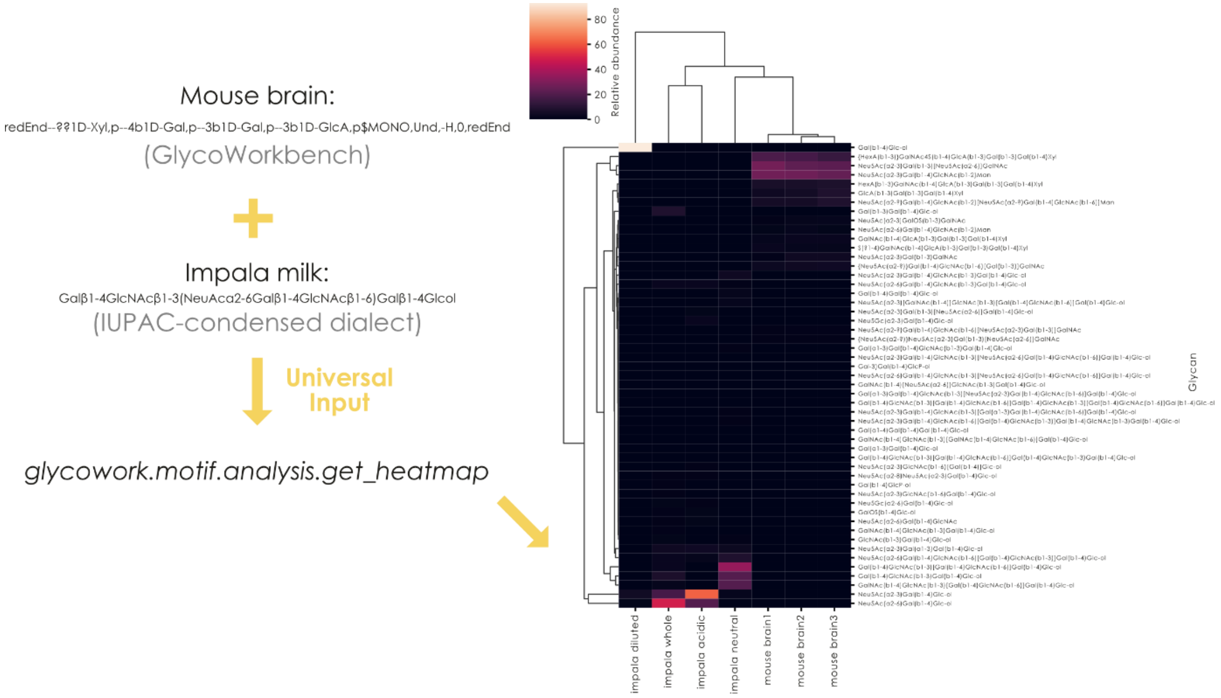


**Supplementary Figure 3. Universal Input supports mixing of nomenclatures.** Using mouse brain *O*-glycans in GlycoWorkbench (GPST000374) and impala free milk oligosaccharides in a dialect of IUPAC-condensed (GPST000317), we present a proof-of-concept experiment of using separate nomenclatures in the same dataset, used for hierarchical clustering via *glycowork.motif.analysis.get_heatmap*.

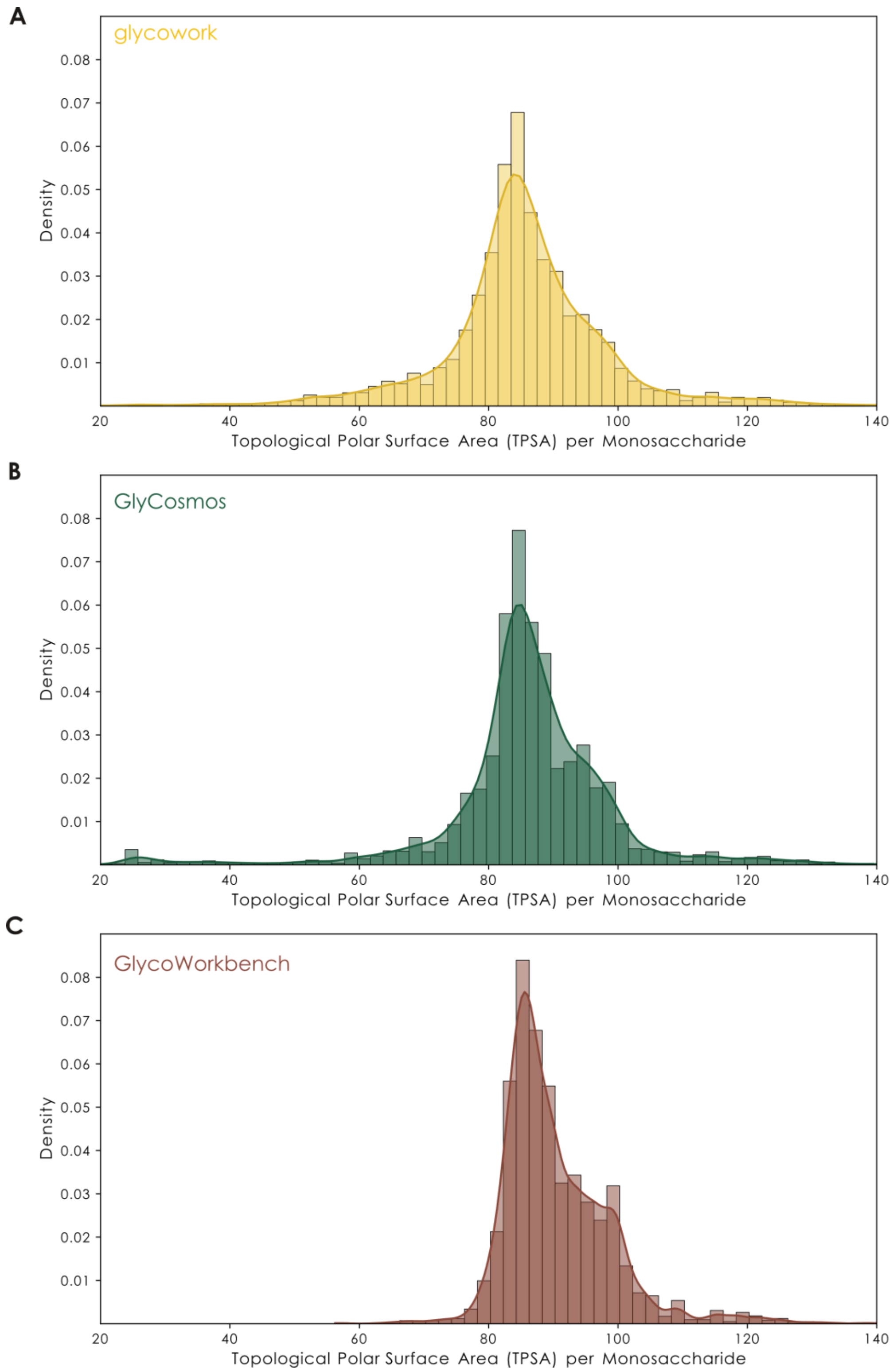


**Supplementary Figure 4. Universal Input enables calculating arbitrary chemical features from fully defined glycans. A-C)** We used 24,626 chemically fully defined glycans from *glycowork.glycan_data.loader.df_glycan* (v1.6.1; IUPAC-condensed), 34,639 chemically fully defined glycans from GlyCosmos (WURCS), and 23,135 chemically fully defined glycans from GlycoWorkbench. Glycan representations from these resources were converted into SMILES string via *canonicalize_iupac* and GlyLES. Then, using rdkit (version 2025.03.2), we calculated the Topological Polar Surface Area (TPSA) of each glycan, and divided the TPSA value by the number of monosaccharides in the glycan.
